# Spontaneous mutation rate in *Saccharomyces cerevisiae* is lower in nutritional and genotypic conditions reducing specific growth rate

**DOI:** 10.64898/2026.01.06.697716

**Authors:** Linda Porri, Celma Mekki, Pinja Salminen, Paula Jouhten

## Abstract

Even in the absence of mutagenic factors, spontaneous errors in DNA replication generate genetic diversity in microbial populations. The spontaneous mutation rate is not immutable but may be conditionally elevated. However, it remains unresolved whether apparently non-stressful nutritional or genotypic conditions affect the spontaneous mutation rate.

Here, we determined the spontaneous mutation rate of haploid *Saccharomyces cerevisiae* CEN.PK113-7D in three nutritional and three genotypic conditions. The nutritional and genotypic conditions influenced the specific growth rate of the *S. cerevisiae* population. Thus, we extended the established fluctuation assay for spontaneous mutation rate determination with *CAN1* as a reporter gene to populations with different generation times. We applied the method to determine the spontaneous mutations rates in wild type *S. cerevisiae* grown on glucose and ammonium, raffinose and ammonium, and glucose and L-tryptophan as sole carbon and nitrogen sources. Alike we determined the spontaneous mutation rates of two engineered *S. cerevisiae* strains (i.e., representing genotypic conditions different from wild type) on glucose and ammonium medium. The engineered strains had two to three heterologous or variant genes integrated into the genome, common to simple heterologous small molecule production host strains. In all alternative nutritional and genotypic conditions, the spontaneous mutation rate of *S. cerevisiae* was reduced compared to wild type growing on glucose and ammonium medium.

Spontaneous mutation rate is fundamentally relevant for evolvability of strains, but it may also influence the performance robustness of microbial populations in applications such as food or beverage fermentation or biotechnological chemical production. Our novel findings are important for biotechnological processes using microbial cells, often engineered and cultivated in unnatural chemical environments.

## INTRODUCTION

Spontaneous mutations arising in DNA replication are a fundamental source of genetic variation, enabling populations of microbial cells to adaptively evolve. The frequency at which spontaneous mutations occur has been found to vary by orders of magnitude between different organisms or strains such as wild type *Saccharomyces cerevisiae* strains ^1-3^. The variation in mutation rates is genotype dependent, largely due to functional diversity of DNA replication and repair systems ^4^ and shows dependence on genome size ^5^. Furthermore, in recent experimental evolution studies, different stressful chemical environments have been observed to elevate the rate of occurrence of spontaneous mutations in *S. cerevisiae* and relatively benign levels of nutrient starvation produce distinct mutational patterns ^6,7^. Stressful chemical environments were indicated by a reduced fitness, represented by population growth rates. In contrast, in *Escherichia coli*, nutrient limitation has been found to induce changes in the spectrum of spontaneous mutations rather than a rise in the overall mutation rate ^8,9^. The breadth of nutritional conditions and underlying mechanisms of the consequent variation in spontaneous mutation rates, remain poorly understood but are of critical interest for both evolutionary biology and biotechnology.

Apart from being genotype and condition dependent, the spontaneous mutation rate is maintained by selection above what would be possible by fidelity mechanisms ^10,11^. The spontaneous mutations underlie evolutionary adaptation, but a too high spontaneous mutation rate hinders rather than mediates adaptation ^12^. This is due to spontaneous mutations including deleterious and neutral mutations in addition to some beneficial (e.g., in yeast estimated up to ∼6% among fitness-affecting) ^13,14^. Fitness-beneficial mutations may have rather opposite effects on other traits, in particular those developed through genetic engineering or selection protocols to overproduce e.g., pleasant fermentation aroma or heterologous compound of commercial value. Thus, the spontaneous mutation rates in genotypic conditions beyond wild types are relevant for industrial application of microbes. Here, we determined the spontaneous mutation rate of haploid *Saccharomyces cerevisiae* CEN.PK113-7D in three nutritional and three genotypic conditions that reduced the specific growth rate. Thus, we extended the established fluctuation assay for spontaneous mutation rate determination with *CAN1* as a reporter gene ^1,15^ to populations with different generation times. We applied the method to determine the spontaneous mutations rates in wild type *S. cerevisiae* grown on glucose and ammonium, raffinose and ammonium, and glucose and L-tryptophan as sole carbon and nitrogen sources. Alike we determined the spontaneous mutation rates of two engineered *S. cerevisiae* strains (i.e., representing genotypic conditions different from wild type) on glucose and ammonium medium. The engineered strains had two to three heterologous or variant genes integrated into the genome, common to simple heterologous small molecule production host strains.

## MATERIALS AND METHODS

### Media and strain construction

For cloning, *E. coli* strains were grown in lysogeny broth (LB) with 100 µg/ml ampicillin (MERCK). For transformation, *S. cerevisiae* was cultivated on YPD containing 20 g/L of bacteriological peptone (Neogen), 10 g/L of yeast extract (Neogen), and 20 g/l D-glucose (VWR Chemicals). For selection of plasmids after and during transformation, YPD medium with 200 µg/ml nourseothricin dihydrogen sulfate (NAT) (Jena Bioscience, AB-101) and 200 µg/ml Geneticin® G-418 sulfate (G418) (MERCK) was used. *S. cerevisiae* transformation was performed using the standard lithium acetate protocol (Gietz, 2014) and the CRISPR/Cas9 protocol of the EasyClone kit (Jessop-Fabre et al., 2016), with expression plasmids linearized with NotI enzyme (FD0596, Thermo Scientific). Correct integration was confirmed with QuickLoad Taq PCR (NEB).

Four chemical environments were used for the accumulation of spontaneous mutations. The composition of the synthetic defined (SD) base medium lacking amino acids or ammonium salts used here was described by Jiang et al., 2022 ^1^. The base medium was supplemented with D-glucose (20 g/L; 0.666 mol C/L) with or without added 10 mM lithium chloride (Glc-NH4 or Glc-NH4-LiCl), or D-raffinose (18.66 g/L; 0.666 mol C/L) and ammonium sulfate (6.7 g/L; 0.0756 mol YAN (yeast assimilable nitrogen)/L; Raf-NH4). Another chemical environment with reduced YAN in a form of a single alternative nitrogen source was prepared by supplementing the base medium with D-glucose and L-tryptophan (7.72 g/L; 0.0378 mol YAN/L; Glc-Trp).

### Strains

The haploid *Saccharomyces cerevisiae* strain CEN.PK113-7D (MATa, *URA3, HIS3, LEU2, TRP1, MAL2-8c, SUC2*), referred to as *wild type* strain, was provided by Euroscarf. The strain referred to as *indigoidine synthetase* (MATalpha, *MAL2-8c SUC2 XII-5::adh1t-sfp<pgk1p-tdh3p>BpsA-eno2t::NatMX*) was engineered previously ^16^ to harbor heterologous indigoidine synthase (BpsA) and a 4′-phosphopantetheinyl transferase (Sfp) ^17^ integrated in the well-characterized genomic locus XII-5 ^18,19^. Similarly, the strain referred to as *tyrosinase* (MATa, *URA3, HIS3, LEU2, TRP1, MAL2-8c, SUC2, XII-5::<tdh3p>melA, X-2::<hhf2p>ARO3*^K222L^*-tpgk1t<tef1p>ARO4*^K229L^-*eno2t*) was engineered to harbor tyrosinase encoding melA from *Rhizobium etli* ^20^ and gene variants *ARO3*^K222L 21^ and *ARO4*^K229L 22^ relieving the allosteric regulation of the encoded enzymes.

### Growth profiles and growth rate estimation

Four clones per strain were transferred from YPG agar (20 g/L of bacteriological peptone (Neogen), 10 g/L of yeast extract (Neogen), 30 g/L D-(+)-galactose (Sigma-Aldrich), and 20 g/L of bacteriological agar (VWR Chemicals)) plates into 3 ml growth medium, incubated at 30 °C and shaking at 220 rpm until an OD600 of 10 was reached. A cell suspension volume of 150 µl, with a starting OD600 of 0.02, was prepared and dispensed into honeycomb plates, providing 10 replicates per strain per medium. Growth profiles were recorded using Bioscreen C (Bioscreen and Growth Curves Ltd., Finland) at 30 °C, high amplitude, high speed and continuous shaking. Turbidity at 600 nm was measured in 15-minute intervals. An average blank value was subtracted from the observed optical density (OD) values, followed by a correction for non-linearity at higher cell densities according to Warringer and Blomberg (2003) ^23^. The corrected OD values were log-transformed to linearize the growth data and to identify the exponential phase. A linear model was fitted to the log-transformed data against time, and the specific growth rate was estimated from the slope of this relationship.

### Mutation rate estimation

Fluctuation assays using L-canavanine sulfate salt and *CAN1* as a reporter gene were conducted following the high-throughput protocol published by Jiang et al. (2022) ^1^. We diverted from the protocol to account for growth rate differences between strains as follows: To minimize confounding effects of population size on mutation rate estimates ^24^, we set a starting population density of 1000 cells in 75 µl medium (13 cells/µl) was prepared for all strains and environments. A minimum of 96 parallel cultures, 24 for total population size and 72 for canavanine-resistant (can^R^) colony detection, for each of three biological replicates was incubated at 30 °C, 220 rpm shaking and humidification of the incubator to reduce evaporation. We optimized the incubation times for all strains in all environments based on their specific growth rates to achieve a similar number of cell divisions during the mutation accumulation period for all, ranging from 24 to 75 h. Total colony counts (*Nt*) were obtained on yeast peptone agar plates with 2% glucose, and can^R^ colony counts (*m*) on SC-Glc-NH4-agar plates with arginine-serine-dropout and L-canavanine sulfate (600 mg/L; C9785, Sigma-Aldrich) supplementation. Data analysis was performed using the rSalvador package v1.7 ^1,25^. The mutation rate (*µ*) of the *CAN1* locus was determined as *µ = m/Nt* canavanine resistance mutations per gene per cell division ^1^.

### Statistical analysis

Differences in specific growth rates and mutation rate estimates, respectively, between strains and conditions were evaluated using One-Way ANOVA and post-hoc Tukey’s range tests. Statistical significance was concluded at an adjusted p-value lower than 0.05. Ninety-five percent confidence intervals (CI) of *m* were calculated using the method described by Lang (2018) ^15^. The 95 % CI of µ were calculated by dividing the lower and upper confidence limits of *m* by *Nt* ^26^.

## RESULTS

### Markedly reduced growth rates upon alternative nutritional and genotypic conditions

*S. cerevisiae* CEN.PK113-7D was grown in three different nutritional conditions with (i) D-glucose and ammonium (Glc-NH4), (ii) D-glucose and L-tryptophan (Glc-Trp), and (iii) D-raffinose and ammonium (Raf-NH4) as the sole carbon and nitrogen sources to determine maximum specific growth rates in each. In addition, the maximum specific growth rate of *S. cerevisiae* exposed to lithium chloride in Glc-NH4 was determined. Lithium chloride is a known environmental stressor previously shown to cause slow growth and increase the spontaneous mutation rate ^7^ but used in lower concentration due to the heightened salt sensitivity of *S. cerevisiae* CEN.PK113-7D ^27^. *S. cerevisiae* CEN.PK113-7D grew in multi-well plate cultures in Bioscreen instrument with the highest apparent maximum specific growth rate of 0.519 ± 0.019 h^-1^ on Glc-NH4. In comparison to growth on Glc-NH4 we observed a decrease to 0.402 ± 0.027 h^-1^ (adj. p = 1.873^-13^; n=10) on Raf-NH4 and to 0.370 ± 0.017 h^-1^ (adj. p = 1.004^-13^; n=10) on Glc-NH4-LiCl. A more than four-fold growth rate reduction was observed when the cells grew on limited YAN and single amino acid as a nitrogen source in Glc-Trp leading to a low maximum specific growth rate of 0.120 ± 0.004 h^-1^ (adj. p = 1.003^-13^; n=10; figure 1A, supplementary table 1). The maximum specific growth rate on Glc-Trp was significantly lower than on Raf-NH4 (adj. p = 1.003^-13^) and on Glc-NH4-LiCl (adj. p = 1.003^-13^), and lower on Glc-NH4-LiCl than on Raf-NH4 (adj. p = 0.004; n=10). Thus, the maximum specific growth rate of *S. cerevisiae* CEN.PK113-7D varied nutritional and stressful conditions between 0.120 ± 0.004 h^-1^ (Glc-Trp) and 0.519 ± 0.019 h^-1^ in (Glc-NH4).

**Figure 1.**
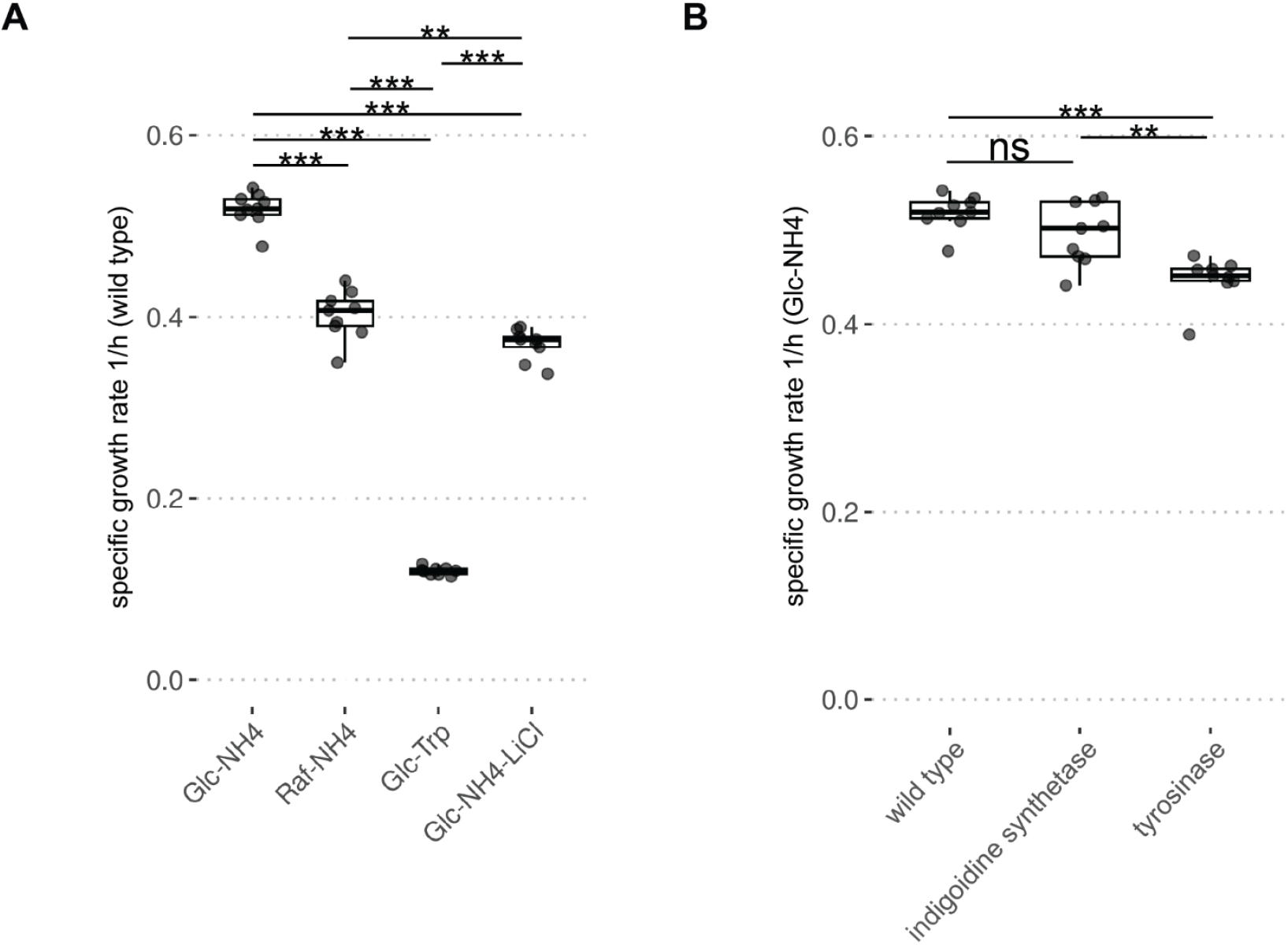
Specific growth rate response to nutritional and genetic conditions. a) The maximum specific growth rates (as averages of ten cultures) of *S. cerevisiae* CEN.PK113-7D in three nutritional conditions: on Glc-NH4 (0.519 ± 0.019 h^-1^), on Raf-NH4 (0.402 ± 0.027 h^-1^) and on Glc-Trp (0.370 ± 0.017 h^-1^) and in presence of a stressor on Glc-NH4-LiCl (0.370 ± 0.017 h^-1^). b) The maximum specific growth rates (as averages of ten cultures) in three genotypic conditions on Glc-NH4: wild type (0.519 ± 0.019 h^-1^) (as in a), *indigoidine synthase* strain (0.496 ± 0.033 h^-1^), and *tyrosinase* strain (0.448 ± 0.024 h^-1^). The differences in the maximum specific growth rates are indicated (ns > 0.05 adj. p; * < 0.05 adj. p; ** < 0.005 adj. p; *** < 0.0005 adj. p; n=10).

Two engineered strains of *S. cerevisiae* CEN.PK113-7D background represented genotypic conditions different from wild type. Integration of indigoidine synthase (BpsA) and a 4′-phosphopantetheinyl transferase (Sfp) ^17^ in the well-characterized genomic locus XII-5 ^18,19^ in *S. cerevisiae* CEN.PK113-7D ^16^ did not reduce the maximum specific growth rate on Glc-NH4 compared to the wild type (i.e., 0.496 ± 0.033 h^-1^; adj. p = 0.172; n=10). This strain is later referred to as *indigoidine synthetase* strain. In contrast, *S. cerevisiae* CEN.PK113-7D having tyrosinase encoding melA from *Rhizobium etli* (Cabrera-Valladares et al. 2006) and gene variants *ARO3*^K222L^ and *ARO4*^K229L^ encoding two isoenzymes of 3-deoxy-D-arabino-heptulosonate-7-phosphate (DAHP) synthase relieved in allosteric regulation of aromatic amino acid biosynthesis grew significantly slower than the wild type, with a maximum specific growth rate of 0.448 ± 0.024 h^-1^ (adj. p = 0.0000155; n=10; figure 1B). This strain is later referred to as *tyrosinase* strain. The *tyrosinase* strain grew also slower than the *indigoidine synthase* strain (adj. p = 0.00164; n=10; figure 1B, supplementary table 1).

### Decreased rates of spontaneous mutations upon alternative nutritional or genotypic conditions

Next, we investigated spontaneous mutation rates of *S. cerevisiae* CEN.PK113-7D in the three nutritional and the three genotypic conditions and in the presence of the stressor lithium chloride. We utilized the high throughput fluctuation assay protocol recording *CAN1* loss-of-function events ^1^. In the fluctuation assay, the spontaneous mutation rates are estimated by first growing multiple independent cultures from small inocula, and then, selecting for loss-of-function mutations in the *CAN1* gene. *CAN1* encodes an arginine permease allowing an uptake of canavanine, a toxic analog of L-arginine. Thus, the loss-of-function mutations in *CAN1* confer resistance to canavanine. To prevent bias in the spontaneous mutation rate estimates due to the differences in the maximum specific growth rates, we synchronized the number of generations across the conditions during the mutation accumulation.

Independent mutation rate estimates as canavanine resistance mutations per gene per cell division from three biological replicates were obtained per strain and environment or genetic condition. Exposure to any altered nutritional condition, whether raffinose as alternative carbon source (7.77^-8^ ± 1.04^-8^ mutations per division; adj. p = 0.002; n=3) or tryptophan as alternative nitrogen source (7.35^-8^ ± 2.16^-8^ mutations per division; adj. p = 0.001; n=3) lead to a decrease in the average rate of spontaneous mutations in comparison to optimal growth conditions (Glc-NH4; figure 2AC; supplementary table 2). In contrast, no change in mutation rate was observed when the strain was exposed to 10 mM lithium chloride (Glc-NH4-LiCl; 2.40^-7^ ± 2.69^-8^ mutations per division; adj. p = 0.234; n=3). No difference was detected between Raf-NH4 and Glc-Trp (adj. p = 0.997; n=3), while mutation rates in both Raf-NH4 (adj. p = 0.002; n=3) and Glc-Trp (adj. p = 0.002; n=3) were markedly lower than in the lithium chloride containing environment (figure 2 AC). A marked decrease in mutation rates was observed for both *indigoidine synthase* (7.65^-8^ ± 1.21^-8^ mutations per division; adj. p = 0.002; n=3) and *tyrosinase* strains (5.15^-8^ ± 2.23^-8^ mutations per division; adj. p = 0.001; n=3) compared to the wild type strain (1.97^-7^ ± 3.49^-8^) grown on Glc-NH4. No difference was observed between the engineered strains (adj.p = 0.479; n=3; figure 2BD, supplementary table 2)

**Figure 2.**
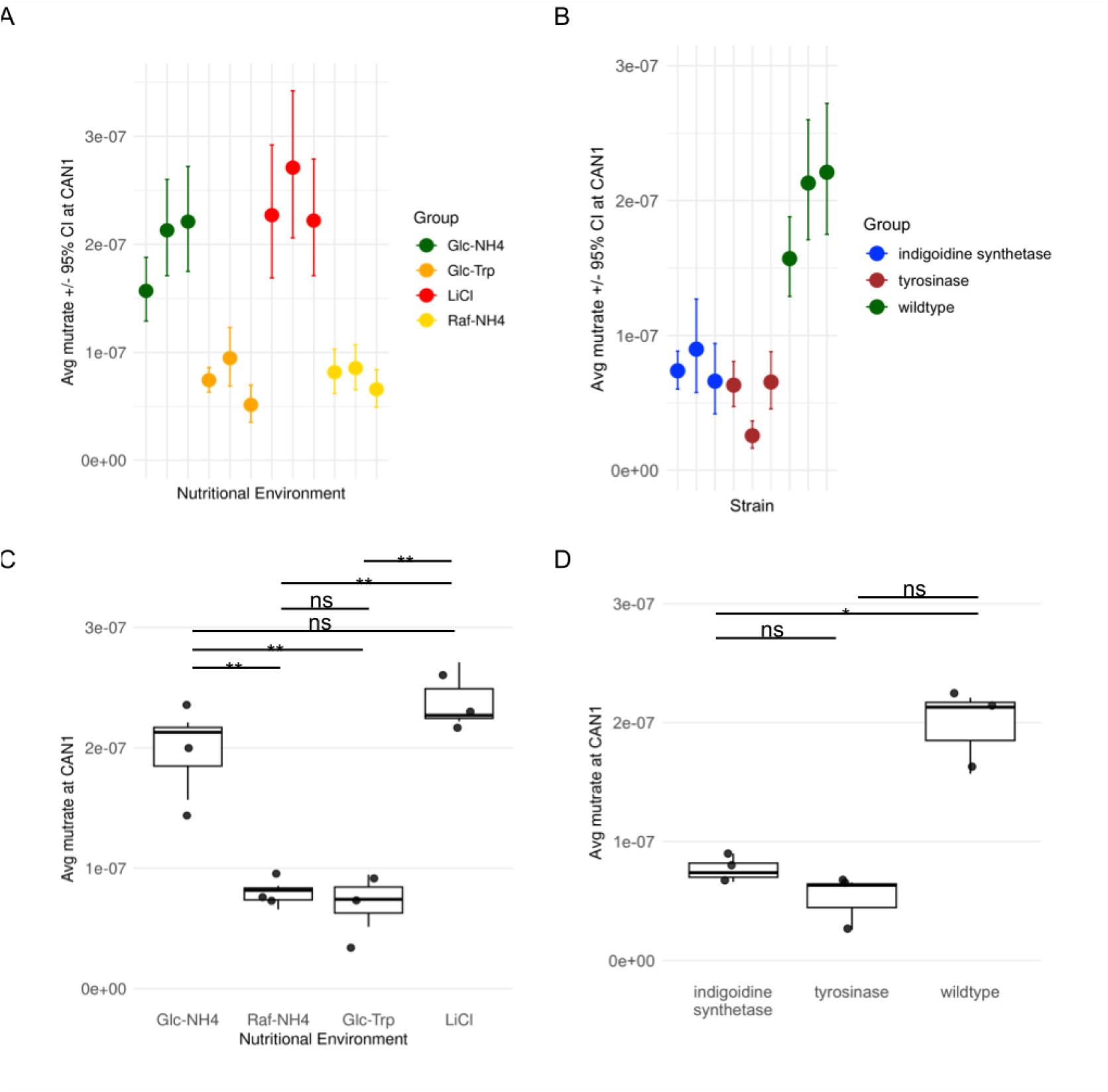
Spontaneous mutation rate estimates decreased upon all alternative nutritional and genotypic conditions but not in response to LiCl exposure. a) Three independent mutation rate (i.e., canavanine resistance mutations per gene per cell division) estimates were obtained per strain and chemical environment, which revealed a decrease upon growth on Raf-NH4 (yellow) and Glc-Trp (orange) compared to growth on Glc-NH4 (green), but a slight increase upon LiCl exposure (red). b) A marked decrease in mutation rates on Glc-NH4 was observed for both *indigoidine synthase* (blue) and *tyrosinase* (brown) strains compared to the wild type strain (green). Dots represent average mutation rate estimates across all parallel cultures of a biological replicate, bars 95% confidence intervals. c) Average spontaneous mutation rates across all biological replicates indicated a statistically significant decrease in Raf-NH4 (7.77^-8^ ± 1.04^-8^ mutations per division) and Glc-Trp (7.35^-8^ ± 2.16^-8^ mutations per division) compared to Glc-NH4 (1.97^-7^ ± 3.49^-8^ mutations per division), and an unchanged rate (2.40^-7^ ± 2.69^-8^ mutations per division) upon LiCl exposure. d) For both *indigoidine synthase* (7.65^-8^ ± 1.21^-8^ mutations per division) and *tyrosinase* (5.15^-8^ ± 2.23^-8^ mutations per division) having strains lower mutation rates were estimated compared to the wild type strain (1.97^-7^ ± 3.49^-8^ mutations per division), but similar ones between themselves (ns > 0.05 adj. p; * < 0.05 adj. p; ** < 0.005 adj. p; *** < 0.0005 adj. p; n=3/).

### Magnitude of growth rate reduction does not translate to a spontaneous mutation rate

The conditions established here, ranging from nutritional and genetic conditions to LiCl exposure, all caused a reduction in the specific growth rate of *S. cerevisiae* CEN.PK113-7D compared to wild type growth on Glc-NH4. However, the spontaneous mutation rate response driven by chemical stress (i.e., lithium chloride) was opposite than by the alternative nutritional or genetic conditions. The growth rate reduction in Raf-NH4 and Glc-Trp nutritional conditions were accompanied with a lower mutation rate than in the common Glc-NH4 environment (figure 3A). Our data suggested that the magnitude of growth reduction does not directly translate to the mutation rate: While the specific growth rate of *S. cerevisiae* CEN.PK113-7D on Glc-Trp was significantly lower than in Raf-NH4 environment (adj. p = 1.003^-13,^; n=10; figure 1A), the mutation rates in the two nutritional environments were similar (adj. p = 0.997; n=3; figure 2AC; figure 3A). Furthermore, distinctly different mutation rates were estimated in Raf-NH4 and Glc-NH4-LiCl conditions (adj. p = 0.002; figure 2AC) despite the only moderately different specific growth rates (0.402 ± 0.027 h^-1^ and 0.370 ± 0.017 h^-1^; adj. p = 0.004; n=10; figure 3A). Genetic conditions of *indigoidine synthase* and *tyrosinase* strains reduced the specific growth rate and reduced the rate of spontaneous mutations compared to the wild type strain (figure 3B). While the *indigoidine synthase* and *tyrosinase* strains’ spontaneous mutation rates were similar (7.65^-8^ ± 1.21^-8^ mutations per division and 5.15^-8^ ± 2.23^-8^ mutations per division, adj.p = 0.479; n=3 figure 2BD; figure 3B), their growth rates were not (0.496 ± 0.033 h-1, 0.448 ± 0.024 h-1; adj. p = 0.00164; n=10; figure 1B; figure 3B).

**Figure 3.**
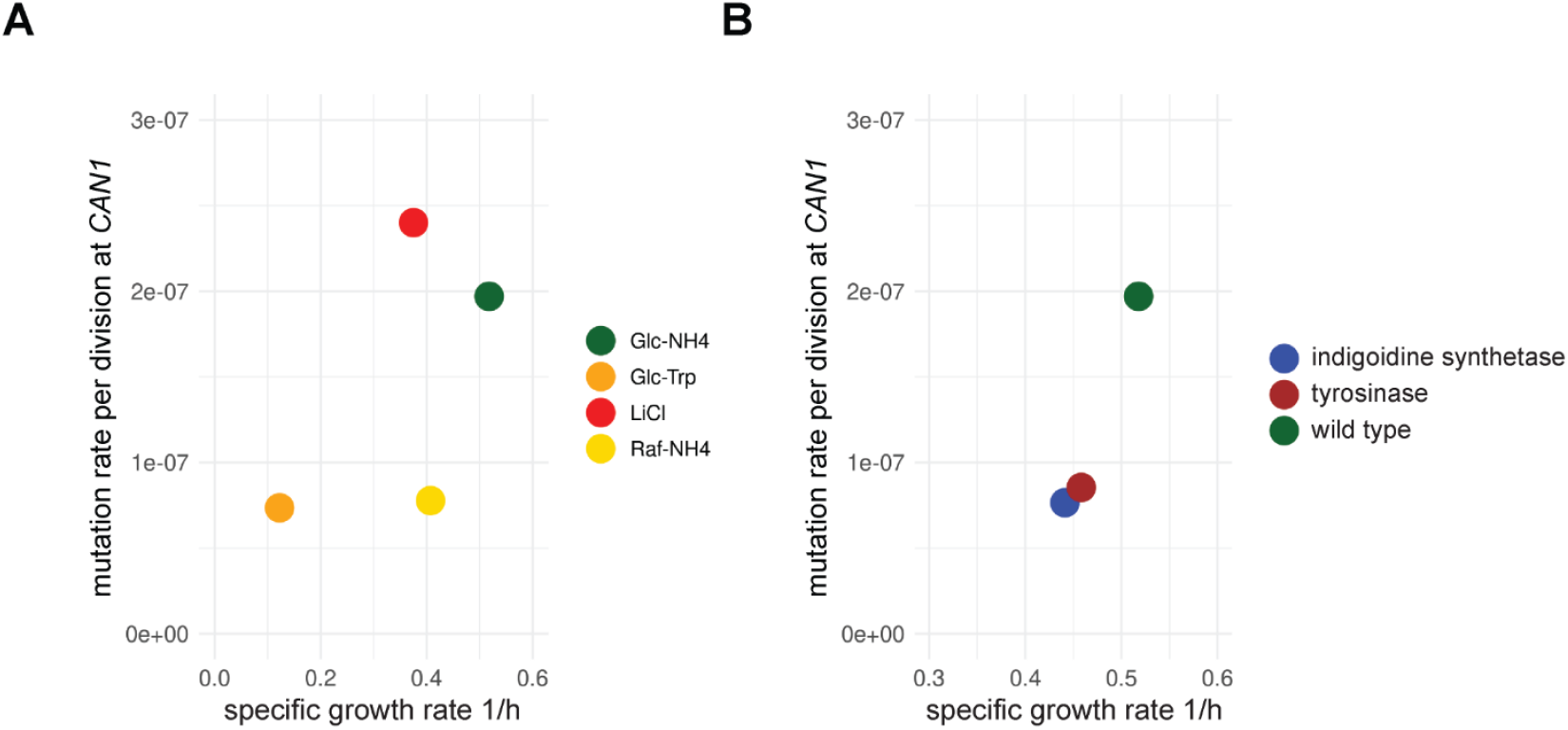
No linear relationship between spontaneous mutation and reduced growth rates. a) Average spontaneous mutation rate per division at *CAN1* and the maximum pecific growth rate (h^-1^) of *S. cerevisiae* CEN.PK113-7D in different nutritional conditions: Glc-Trp (orange), Raf-NH4 (yellow), Glc-NH4 (green), and in the presence of a stressor: Glc-NH4 supplemented with 10 mM LiCl (red). b) Average spontaneous mutation rate per division at *CAN1* and the maximum specific growth rate (h^-1^) of *S. cerevisiae* CEN.PK113-7D in different genetic conditions: wild type, and *indigoidine synthase* and *tyrosinase* strains.

## DISCUSSION

Inspired by low growth rate, stressful, chemical environments having been found to elevate spontaneous mutation rates in *S. cerevisiae* ^7^, we determined the spontaneous mutation rates in alternative nutritional and genotypic conditions, all affecting the specific growth rate. Despite triggering growth rate reductions, neither alternative carbon and nitrogen sources nor the genome editing resulted in increased mutation rates per division, rather opposite.

In all alternative conditions in an absence of stressors, the spontaneous mutation rates of *S. cerevisiae* were reduced. The apparent mutation rates may become reduced due to reasons reducing the cell internal stress. For instance, the spontaneous mutation rate can be reduced due to physiological changes dependent on high cell concentration in a phenomenon called density-associated mutation-rate plasticity (DAMP) characterized in *E. coli* ^24^. DAMP can arise from an enhanced ability to detoxify oxidizing agents (external and internally generated) in a dense population ^28^. However, DAMP was likely not involved in the differences between the mutation rates we estimated since we adjusted the starting population density, the effects of evaporation, as well as the number of generations passed across all mutation accumulation experiments. The differences could still have involved oxidizing agents. For instance, related to our Raf-NH4 conditions, caloric restriction has been shown to reduce ROS released/O_2_ consumed ^29^. The major source of ROS is oxidative phosphorylation, which was repressed in the rest of the conditions, due to excess glucose. However, most stress responsive genes show also growth rate dependent expression ^30^ and slow-growing cells have been found more stress-resistant (Elliot and Futcher, 1993; Slattery and Heidemann, 2007). Apparently in nutritional and genotypic conditions leading to lowered growth rate, maintaining replication fidelity and avoiding deleterious mutations has been an evolutionarily beneficial allocation of cellular resources.

In conclusion, our data supported the idea that slower growth rates in an absence of stressors reduce spontaneous mutation rates. The underlying mechanisms, potentially involving the altered cellular physiology, the total metabolic flux and respiratory activity, remain to be fully elucidated and warrant further investigation across more diverse conditions.

## CODE AVAILABILITY

The code is available at https://version.aalto.fi/gitlab/microbial-physiology/adaco_fluc_assay.

## AUTHOR CONTRIBUTIONS

Linda Porri: Formal analysis, Investigation, Methodology, Visualization, Writing – original draft, Writing - review & editing; Celma Mekki: Investigation, Writing - review & editing; Pinja Salminen: Investigation, Writing - review & editing; Paula Jouhten: Conceptualization, Funding acquisition, Supervision, Formal analysis, Investigation, Writing – original draft, Writing - review & editing.

## ACKNOWLEDGEMENTS

Paula Jouhten acknowledges funding from Novo Nordisk Foundation (NNF22OC0080180).

## SUPPLEMENTARY MATERIAL

**Supplementary table 1**. Maximum specific growth rates of *S. cerevisiae* CEN.PK113-7D in nutritional (Glc-NH4, Raf-NH4, Glc-Trp) and genotypic (wild type, *indigoidine synthase, tyrosinase*) conditions, and in the presence of a stressor (Glc-NH4-LiCl).

**Supplementary table 2**. Spontaneous mutation rates of *S. cerevisiae* CEN.PK113-7D in nutritional (Glc-NH4, Raf-NH4, Glc-Trp) and genotypic (wild type, *indigoidine synthase, tyrosinase*) conditions, and in the presence of a stressor (Glc-NH4-LiCl).

